# Neural mechanisms underlying release-mode-specific abnormalities in dopamine neural activity in a schizophrenia-like model

**DOI:** 10.1101/2023.05.09.540082

**Authors:** Hidekazu Sotoyama

**Affiliations:** Department of Physiology, Niigata University School of Medicine; Department of Molecular Neurobiology, Brain Research Institute, Niigata University

## Abstract

Abnormalities in dopamine function might be related to psychiatric disorders such as schizophrenia. Even at the same concentration, dopamine exerts opposite effects on information processing in the prefrontal cortex depending on independent dopamine release modes known as tonic and phasic releases. This duality of dopamine prevents a blanket interpretation of the implications of dopamine abnormalities for diseases on the basis of absolute dopamine levels. Moreover, the mechanisms underlying the mode-specific dopamine abnormalities are not clearly understood. Here, we demonstrate that the two modes of dopamine release in the prefrontal cortex of a schizophrenia-like model are disrupted by different mechanisms. In the schizophrenia-like model established by perinatal exposure to inflammatory cytokine, epidermal growth factor, tonic release was enhanced and phasic release was decreased in the prefrontal cortex. We examined the activity of dopamine neurons in the ventral tegmental area (VTA), which sends dopamine projections to the prefrontal cortex, under anesthesia. The activation of VTA dopamine neurons during excitatory stimulation (local application of glutamate or NMDA), which is associated with phasic activity, was blunt in this model. Dopaminergic neuronal activity in the resting state related to tonic release was increased by disinhibition of the dopamine neurons due to the impairment of 5HT2 (5HT2A) receptor-regulated GABAergic inputs. Moreover, chronic administration of risperidone ameliorated this disinhibition of dopaminergic neurons. These results provide an idea about the mechanism of dopamine disturbance in schizophrenia and may be informative in explaining the effects of atypical antipsychotics as distinct from those of typical drugs.

**Significance:** I discovered that the hypo-NMDA function occurs in midbrain dopaminergic neurons of a schizophrenia-like model instead of the cerebral cortex, which has been the focus of attention so far. This suggests that the schizophrenia glutamate hypothesis may interact with the dopamine hypothesis. Furthermore, it was elucidated that a subpopulation of dorsal raphe serotonergic neurons inhibits VTA dopaminergic neurons in the resting state, resulting in promotion of social behavior. 5HT2 receptor-mediated regulation of inhibitory inputs to the dopaminergic neurons underlies this serotonergic regulation. In the schizophrenia-like model, this regulation by 5HT2 receptors is impaired. Chronic administration of an atypical antipsychotic ameliorates this abnormality. Therefore, this result may represent a mechanism underlying the differential efficacy between atypical and typical antipsychotics.

## Introduction

Dopaminergic pathways originating from the ventral tegmental area, including prefrontal projections and nucleus accumbens projections, are closely associated with social behavior regulation and cognitive functions [1–5]. It is believed that abnormalities in these dopamine pathways are associated with the pathophysiology of psychiatric disorders such as schizophrenia [6–8]. However, information on these dopaminergic abnormalities still remains controversial with often contradictory findings [9–12]. Against this issue, it has been reported that continuously sustained dopamine level and a transiently fluctuating dopamine level exert opposite effects on information processing in the prefrontal cortex even under an identical concentration of dopamine, implying that actual dopamine signal cannot be determined simply by evaluation of extracellular dopamine levels without consideration of dopamine dynamics [13]. Nevertheless, previous pathophysiological studies of psychiatric disorders have not argued this dual role of dopamine between transient and sustained releases with sufficient discussion.

Previous studies have often argued about psychostimulant-induced dopamine release for abnormal dopamine dynamics in patients with schizophrenia and animal models of schizophrenia because responses to psychostimulants are sensitized in patients with schizophrenia [15–18]. However, changes in dopamine levels underlying the physiological regulation of behaviors differ significantly from this drug-induced dopamine efflux [14,19]. Under the physiological conditions, two independent modes of releases, tonic release and phasic releases, play distinct roles in dopamine transmission. Tonic release contributes to maintaining basal dopamine level in the resting state, and phasic release contributes to behavioral stimulus-triggered transient alteration of dopamine level [20,21]. Therefore, in order to understand the influences of dopamine on psychiatric disorders given the dual role of dopamine corresponding to acute and chronic alterations of dopamine level, it is informative to understand how these two modes of dopamine release are disrupted in these diseases.

To proceed this argument, I have investigated the following animal model with schizophrenia endophenotypes. In hypotheses for etiology of schizophrenia, viral infection in the perinatal period is receiving research attention as one of etiological risk factors for schizophrenia. Several studies have suggested that inflammation caused by viral infection results in induction of epidermal growth factor (EGF) [51, 52, 61–63]. Interestingly, a recent study reported that EGF concentrations are elevated in the blood and cord blood of pregnant women infected with the novel coronavirus SARS-CoV-2 [22]. This finding suggests that viral infection in the perinatal period increases the EGF level in the blood of fetuses/newborns. The immature blood–brain barrier of the fetuses/newborns enables high concentrations of EGF to enter the brain from the blood, causing the impairment of brain development with schizophrenia endophenotypes as we have reported previously [23]. Moreover, a postmortem brain analysis of patients with schizophrenia indicates that EGF receptor (ErbB1) level is elevated in patients with schizophrenia [24]. On the basis of these rationales in the epidemiology and pathology of schizophrenia, we established an animal model in which chronic EGF administration in the perinatal period results in disturbances that are observed in schizophrenia, such as reduced social interaction, impairment in sensorimotor gating, impaired latent inhibition of learning, and deficits in mismatch negativity [23,25,26]. This animal model exhibits abnormal dopamine releases in the prefrontal cortex [27], supporting insights led by some positron emission tomography (PET) studies having investigated prefrontal dopamine in patients with schizophrenia [53–57]. Using this schizophrenia-like animal model, I aimed to elucidate the mechanisms underlying abnormal dopamine releases during resting state and behavioral activation that reflect tonic and phasic dopamine releases.

## Materials and Methods

### Animals

Male newborn Sprague-Dawley rats (SLC Ltd., Hamamatsu, Japan) were housed with a dam under a 12-hr light/dark cycle (light on 8:00 a.m.) in a plastic cage (276×445×205 mm). After weaning [postnatal day (PND) 20–30], rats were separated and housed with 2–3 rats per cage. Each adult rat (PND 56–90) was used in each experiment. Male knockin rats expressing Cre recombinase by a tryptophan hydroxylase 2 gene promoter driving (HsdSage: LE-*Tph2^em1(T2A-Cre)sage^*) (Origin: SAGE Labs, Inc., St. Louis, MO, USA) were also housed under the same conditions. The rats were allowed free access to food and water. All of animal experiments described were approved by the Animal Use and care Committee guidelines of Niigata University and performed in accordance with the guidelines of NIH-USA. Every effort was made to minimized the discomfort of the animals in addition to the number of animals used in the experiments.

### Epidermal growth factor (EGF) treatment

Recombinant human EGF (Higeta Shoyu Co., Chiba, Japan) was dissolved in saline. EGF (0.875 μg/g) was administered subcutaneously (s.c.) each day to half of littermates during PND 2–10. Control littermates received an injection of saline on the same schedule. The dose of EGF used in this study did not produce any apparent growth retardation in rats.

### Drug treatment

Some of animals received risperidone (1.0 mg/kg/day, i.p.; Rispadal®; Janssen Pharmaceuticals Inc., Tokyo, Japan), ritanserin (2 mg/kg/day i.p.; Sigma-Aldrich, St. Louis, MO, USA), or haloperidol (0.6 mg/kg/day; Sigma-Aldrich) for 10–14 days prior to the following electrophysiological tests.

### In vivo microdialysis

The rat was anesthetized using the mixture of midazolam (2.0 mg/kg i.p.; Dormicum; Astellas Pharma Inc., Tokyo, Japan), medetomidine (0.38 mg/kg i.p.; Domitol; Nippon Zenyaku Kogyo. Co, Fukushima, Japan) and butorphanol (2.5 mg/kg i.p.; Vetorphale, Meiji Seika Pharma Co., Tokyo, Japan) and mounted on a stereotaxic apparatus. The skull was exposed, and a hole was drilled for unilateral implantation of a guide cannula (AG-8, Eicom, Kyoto, Japan) into the medial prefrontal cortex (stereotaxic coordinates: 3.2 mm anterior to the bregma, 0.8 mm lateral from the midline, 1.7 mm below from the dura surface), according to Paxions and Franklin [28]. After recovery for 10 days, a microdialysis probes (3-mm active area: A-I-8, Eicom) was inserted into the guide cannula. *In vivo* microdialysis was performed according to the previous study [29]. In the combination with social stimulation, a perfusate of artificial cerebrospinal fluid (147 mM NaCl, 2.7 mM KCl, 1.2 mM CaCl_2_, 0.5 mM MgCl_2_, pH 7) was delivered at 2 μL/min. In the combination with depolarizing stimulation, a perfusate of artificial cerebrospinal fluid (147 mM NaCl, 2.7 mM KCl, 1.2 mM CaCl_2_, 0.5 mM MgCl_2_, pH 7) was delivered at 0.7 μL/min. The perfusion medium was switched to the medium containing a high concentration of potassium (80 mM KCl, 69.7 mM NaCl, 1.2 mM CaCl2, 0.5 mM MgCl2; pH 7) for 60 min (for two fractions). The perfusion medium was then switched back to the original medium. Dopamine in the dialysates was determined using HPLC with electrochemical detection. The mobile phase (48 mM citric acid, 24 mM sodium acetate, 10 mM NaCl, 0.5 mM EDTA, 3 mM sodium dodecyl sulfate, and 15% acetonitrile, pH 4.8) was delivered at 50 μL/min. Dopamine was separated using an analytical column (BDS Hypersil C18 1 × 100 mm; Thermo Fisher Scientific, Yokohama, Japan) and detected using a 3-mm glass carbon electrode given +550 mV (Unijet flow cell, Bioanalytical Systems Inc., West Lafayette, IN, USA). Data analysis was performed using Epsilon LC analysis software (Bioanalytical Systems Inc.). Data were not standardized with the recovery rate.

### In vivo extracellular single-unit recording under anesthetized conditions

Extracellular single-unit recording was performed under chloral hydrate anesthesia (400 mg/kg, i.p.). Anesthetized rats were mounted on a stereotaxic apparatus, and their body temperature was continuously controlled to 37 ± 0.5 °C. The skull overlying the target areas was removed. A glass microelectrode filled with 0.5 M NaCl containing 2 % Pontamine Sky Blue (resistance 10-20 MΛ) was inserted into the ventral tegmental area (VTA). The stereotaxic coordinates were VTA (5.2–5.6 mm posterior (AP) and 0.6–1.0 mm lateral (ML) from the bregma, 7.5–8.5 mm ventral (DV) from the dura; the dorsal raphe (AP −7.5 to −8.0, ML 8.0 to 1.2 with a +10° angle toward the lateral direction, DV −4.8 to −6.5) according to Paxinos and Watson [28]. Each dopamine neuron was recorded for 2 min after 2 min stabilization period. After recording, Potamine Sky Blue was iontophoretically injected (−20 μA for 15 min). The brain was removed and fixed in 4 % paraformaldehyde. To determine the recording position, brain slices (100 μm-thickness) were prepared. By measuring a distances from the dye-injection site, recording positions were determined. Dopaminergic neurons were distinguished by established criteria [typical triphasic action potential with rather longer duration (action potential width from an onset to a negative peak through > 1.1 m sec); a slow firing rate (0.5-10 Hz) with occasional bursting activities] [30,31]. Dopaminergic neuron firing was analyzed with respect to the mean firing rate and spike number / minute because there were no significant differences in burst firing patterns under anesthesia as well as during activation of social behavior between controls and EGF-treated rats in my previous report [27]. I distinguished the activities of putative serotonergic neurons in the dorsal raphe as described previously [32]: (1) the typical biphasic action potential with a marked negative deflection, (2) the characteristic duration (0.2–4.0 ms) with the action potential width from start to the negative trough >1.1 ms, and (3) a regular rhythmic spiking pattern of activity.

### Iontophoresis

During in vivo recording, glutamate, ritanserin, or MDL100907 was introduced by iontophoresis in some experiments. Two barrels of a double barrel glass microelectrode were allocated to unit recording of neural activity (Recording barrel) and introduction of each chemical agent (Introduction barrel) respectively. The introduction barrel was filled with Glutamate solution (50 mM glutamic acid, 10 mM NaCl, pH 8.2), Ritanserin solution (2.5 mM ritanserin, 100 m tartaric acid, 10 mM NaCl, 5% DMSO, pH 2.5), or MDL100907 solution (200 μM MDL100907, 100 m tartaric acid, 10 mM NaCl, pH 4). In the experiment of glutamate introduction, positive current (+4 nA) was applied to the Introduction barrel for preventing diffusion of glutamate before stimulation, and then applied current was switched to negative (−4 nA) to introduce glutamate. In the experiments of introduction of ritanserin and MDL100907, negative current (−4 nA) was applied to the Introduction barrel before the introduction, and then applied current was switched to positive (+4 nA) to introduce these two chemicals.

### In vitro extracellular single-unit recording

*In vitro* single unit recording was performed according to the previous study [33]. The rats were anesthetized with isoflurane, and the brains were quickly removed and cooled for 5 min in an ice-cold slicing solution containing 120 mM choline chloride, 1.25 mM NaH2PO4, 2.5 mM KCl, 7 mM MgCl2, 0.5 mM CaCl2, 26 mM NaHCO3, 15 mM glucose, 1.3 mM ascorbic acid, 1 mM sodium pyruvate, and bubbled with 95% O2 and 5% CO2. Each brain was mounted on a vibratome (Pro7; Dosaka EM Ltd., Kyoto, Japan) and cut to 400 μm thick horizontal slices in the same slicing solution. Typically, two to three hemispheral slices were obtained from one brain. The slices were incubated at 34°C for 30 min in a Krebs solution containing 124 mM NaCl, 1.0 mM NaH2PO4, 3.0 mM KCl, 1.2 mM MgCl2, 2.4 mM CaCl2, 26 mM NaHCO3, 10 mM glucose, 1.0 mM ascorbic acid, 1 mM sodium pyruvate. Following incubation at 25°C for more than 30 min, a VTA slice was transferred to a recording chamber where Krebs solution was perfused. In some of the experiments, the perfusion solution was switched to that supplemented with 10 μM bicuculline plus 6-cyano-7-nitroquinoxaline-2,3-dione (CNQX; 10 μM), (plus) (2R)-amino5-phosphonopentanoate (APV; 50 μM) (all from Sigma-Aldrich). At N-methyl-D-aspartic acid (NMDA) stimulation, the perfusion solution was switched to Krebs solution containing NMDA (20 μM). At serotonin stimulation, the perfusion solution was switched to Krebs solution containing serotonin (100 μM). At 5HT2 receptor blockade, the perfusion solution containing ritanserin (10 μM) was used. A glass microelectrode filled with 2.0 M NaCl was inserted into a lateral area of the VTA at the same coordinates of the in vivo extracellular recording. Neuronal signals were amplified using an amplifier (MEZ-8301; Nihon Kohden) connected to a high gain amplifier (AVH-11; Nihon Kohden). The signals were displayed on an oscilloscope (VC-11; Nihon Kohden), and transferred via a digitizer (Digidata 1200; Molecular Devices) to a computer equipped with recording software (Clampex 7, Molecular Devices).

### Injection of AAV vectors

The rat was anesthetized using the mixture of midazolam (2.0 mg/kg i.p.; Dormicum; Astellas Pharma Inc., Tokyo, Japan), medetomidine (0.38 mg/kg i.p.; Domitol; Nippon Zenyaku Kogyo. Co, Fukushima, Japan) and butorphanol (2.5 mg/kg i.p.; Vetorphale, Meiji Seika Pharma Co., Tokyo, Japan). The rat was mounted on a stereotaxic apparatus. The rat was injected with 1.0 μl per site of rAAV5-hsyn-DIO-hM3Dq-mCherry, or rAAV5-hsyn-DIO-mCherry (around 5.0 x 10^12^ vg/ml for each) (Addgene, Watertown, MA, USA) in the dorsal raphe nucleus [Anterior-posterior (AP): -7.5 mm; Lateral (L): +0.8 mm (+10° angle toward the lateral direction); Dorsal-ventral (DV): -6.0 mm] at 0.1 μl/min using a 31-gauge Hamilton syringe. After at least 14 days, the rat was used for each experiment.

### Treatment of Clozapine N-oxide

Clozapine N-oxide (CNO) (Hello Bio Ltd. Bristol, UK) was dissolved in mixture of dimethyl sulfoxide (DMSO) and 20mM citric acid, and then diluted with saline (the final concentration is 50 μg/μl). For the chronic treatment, the CNO solution was filled in a mini-osmotic pump (Alzet model 2002, Durect Co., CA, USA), and then the mini-osmotic pump was implanted under the skin. The dose of CNO was determined according to our previous study [13].

### Social interaction test

The index for social interaction of rats was measured according to a previous stud [13]. A test rat (12 - 16 weeks old) was placed in the open field box (45 cm length x 45 cm width x 30 cm height, MED Associates) under a moderate light level (200 Lx). After 60 min habituation period, a stranger rat was placed in the open field box. And social interactions were monitored for 10 min. Scoring of social interaction times and duration was based on sniffing behavior, defined as active chasing of the partner, shaking the nose close to the partner, contact the partner with the nose (Partners were same age). All tests were videotaped and scored in a blinded fashion with the aid of the video tracking software Etho-Vision XT (Noldus; Wageningen, the Netherlands).

### Statistical analysis

To compare the differences between two groups, Mann–Whitney’s *U* test was used. To compare mean firing rates of VTA dopamine neurons between four or five groups, Kruskal–Wallis’s test was initially used, and followed by Scheffe’s multiple comparison as *post-hoc* comparisons. Dopamine concentrations in dialysates and time-course of normalized or non-normalized spike numbers were initially analyzed using repeated-measures ANOVA with group as a between-subject factor and time as a within-subject factor, and followed by Fisher’ LSD multiple comparison for *post-hoc* comparisons. Correlations were determined by Pearson’s correlation coefficient. *P* < 0.05 indicated statistical significance. Statistical analyses were performed using SPSS software (SPSS, Yokohama, Japan).

## Results

We have previously reported that rats treated with EGF in the perinatal period (postnatal days 2–10) (Figure 1A) exhibit schizophrenia-like behavioral abnormalities such as impaired social behavior after maturity (after 8 weeks of age) [23]. We demonstrated the induction of extracellular dopamine levels by social stimulation in the prefrontal cortex of adult EGF-treated rats (Figure 1B). Resting state dopamine levels were elevated in EGF-treated rats [repeated ANOVA: F(1,18) = 4.58, P = 0.046]. Conversely, changes in dopamine levels induced by social stimulation were reduced in EGF-treated rats [repeated ANOVA: F(1,18) = 5.34, P = 0.032]. To explore whether the changes in dopamine levels reflect neural activity, we investigated the alterations in dopamine levels induced by depolarizing stimulation (Figure 1C). We observed no significant difference in the changes in dopamine levels induced by high potassium stimulation between controls and EGF-treated rats [repeated ANOVA: F(1,12) = 4.45, P = 0.056]. This result suggests that the amount of dopamine that can be released by depolarization is not altered in EGF-treated rats, so that the decrease in social-stimulation-induced elevation in dopamine levels is not originated from a decrease in the dopamine content packaged in synaptic vesicles. Moreover, because dopamine levels after high potassium stimulation show no interaction with time [repeated ANOVA: F(1,12) = 4.45, P = 0.423], there may be no difference in dopamine reuptake between EGF-treated rats and controls. The increase in basal dopamine level is reported to enhance activation of presynaptic dopamine autoreceptors, resulting in reduction of stimulus-induced phasic dopamine release [21]. However, dopamine neurons projecting to the prefrontal cortex lack Girk2-coupled dopamine D2 autoreceptors [31]. Therefore, the blunted phasic dopamine release in the prefrontal cortex of EGF-treated rats is unlikely to be the consequence of presynaptic inhibition by autopreceptors. The changes in dopamine levels in the resting state and in response to social stimulation can be assumed to be largely affected by the activity of dopamine neurons at each stage.

**Figure 1.**
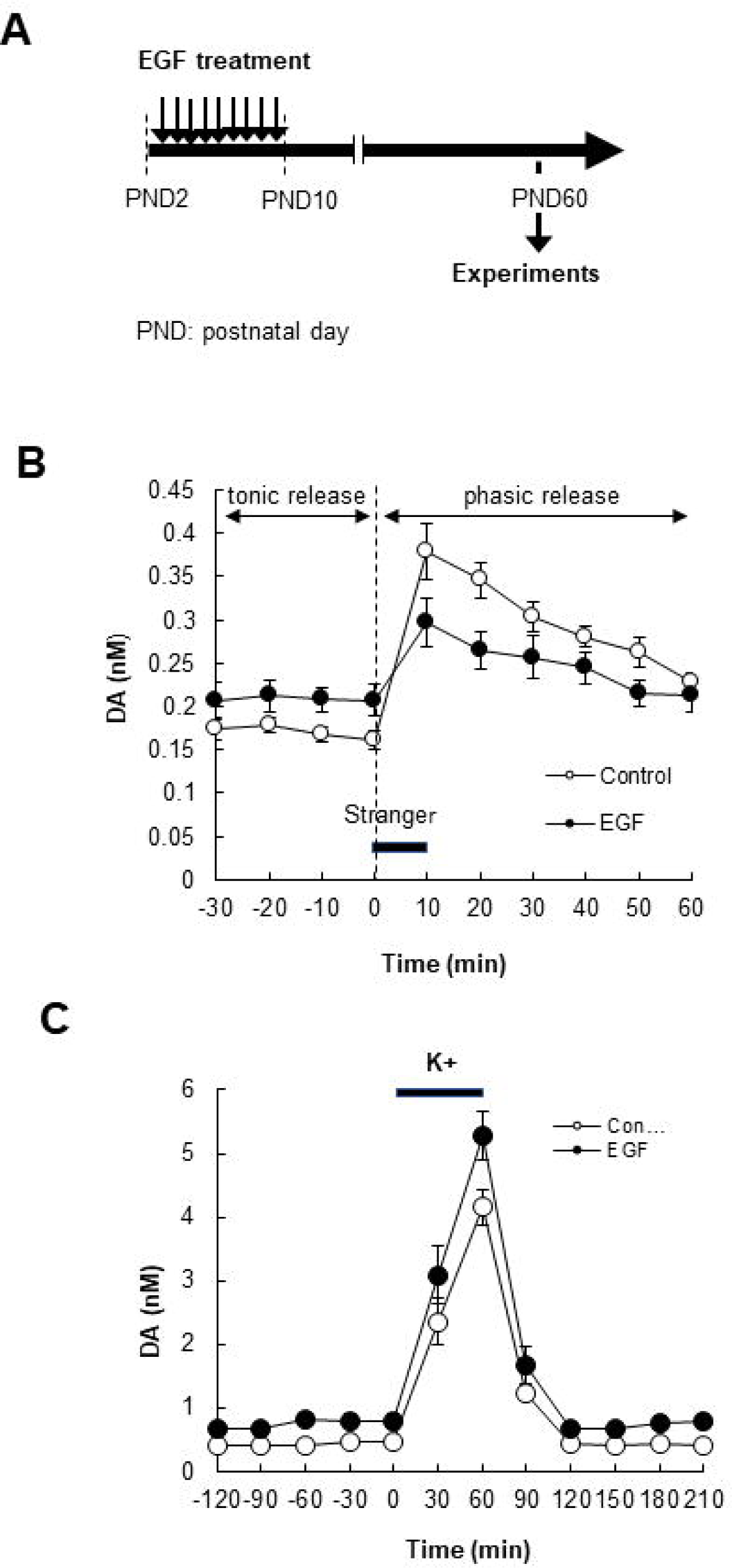
Abnormalities in tonic and phasic dopamine releases in the prefrontal cortex of EGF-treated rats. (A) Schematic diagram of EGF treatment and experiments. EGF was administered daily during postnatal day (PND) 2–10. Microdialysis experiments were performed over PND60 in the prefrontal cortex. (B) Social stimulus evoked dopamine efflux. After monitoring dopamine release in the resting state (tonic release), the EGF-treated rats or controls were encountered to a stranger male rat for 10 min as a social stimulus. The social stimulus increased the dopamine level (phasic release). (N = 10 for each). (C) Dopamine efflux induced by high K+ stimulation was monitored. High K+ was infused from the microdialysis probe for 60 min (N = 7 for each).

Previously, I reported that VTA dopamine neurons in EGF-treated rats exhibit higher firing activity in the resting-state under awake condition compared with controls [27]. Furthermore, I showed that social stimulus increased firing activity to the same level in controls as those of EGF-treated rats, while the social stimulus did not increase further from the high firing in EGF-treated rats [27]. From these results, I thought that EGF-treated rats may have blunt responsiveness to inputs during behavioral activation. Several studies have reported that excitatory inputs from the outside to the VTA are required to induce phasic dopamine release [34,35]. Therefore, I hypothesized that the responsiveness of VTA dopaminergic neurons to excitatory inputs is reduced in EGF-treated rats. To explore this hypothesis, I investigated the changes in VTA dopaminergic activity in vivo when glutamate was applied during the in vivo recording of neural activity (Figure 2A). The administration of glutamate at the recording site using iontophoresis induced a trend of blunt activation in EGF-treated rats compared to that in controls [not statistically significant] (Figure.2B). Normalization of this result revealed that the response of VTA dopamine neurons to glutamate is significantly blunt in EGF-treated rats [repeated ANOVA: F(1,27) = 4.53, P = 0.043] (Figure 2C). Furthermore, using VTA slices, we examined the changes in VTA dopaminergic activity when NMDA was applied. Similar to the result illustrated in Figure 2C, NMDA induced only a smaller hyperactivity of VTA dopaminergic neurons in EGF-treated rats than in controls [repeated ANOVA: F(1,25) = 7.35, P = 0.012] (Figure 3). Therefore, EGF may reduce the NMDA-receptor-mediated regulation of VTA dopaminergic neuronal activity, which may cause a failure to respond to stimulus-induced inputs to VTA dopaminergic neurons. This may contribute to the reduction of the phasic dopamine release in the prefrontal cortex of EGF-treated rats.

**Figure 2.**
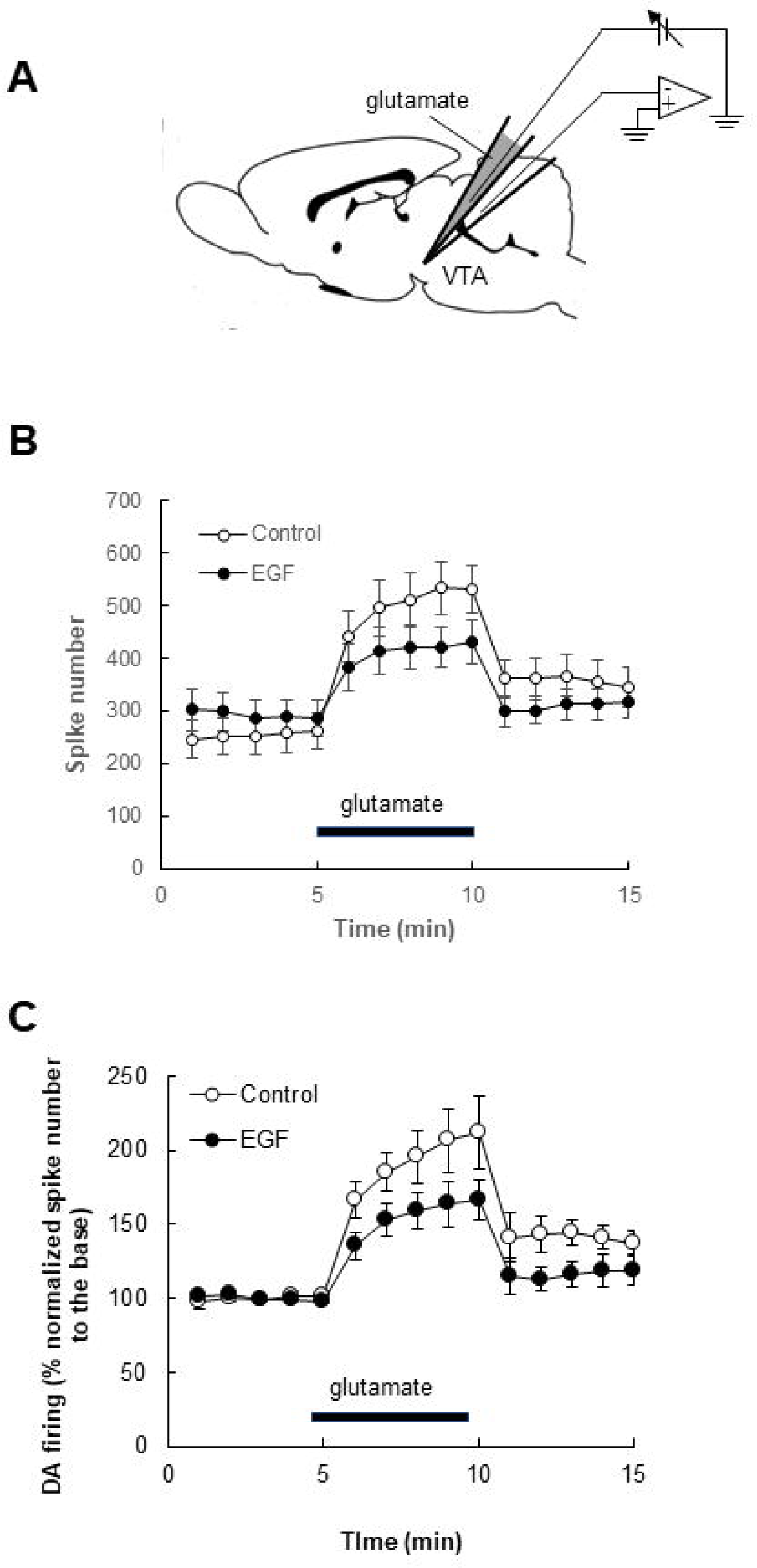
VTA dopaminergic neurons exhibited blunt response to glutamatergic stimulation in EGF-treated rats during in vivo recording. (A) Glutamate was introduced into the single unit recording site by iontophoresis under anesthetized condition. (B) Non-normalized results of responses of VTA dopamine neurons to glutamate application. (C) Normalization of the spike numbers to the base shows that glutamate-induced activation of VTA dopaminergic neurons in EGF-treated rats was blunt compared with controls (N = 12 cells for controls, and N = 18 cells for EGF-treated rats; Animal: N = 4–5).

**Figure 3.**
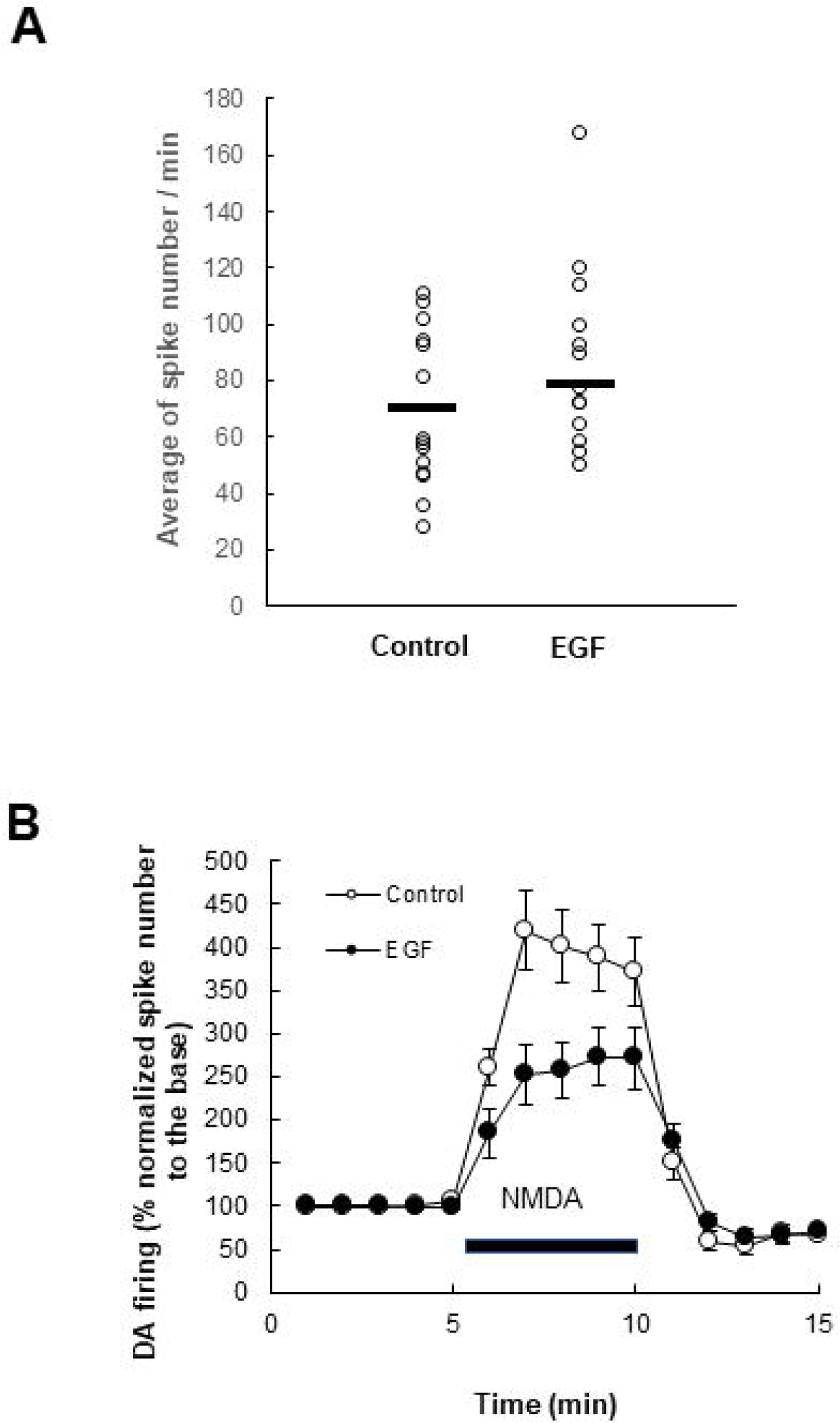
Direct NMDA-receptor-mediated activation of VTA dopaminergic neurons is blunt in EGF-treated rats compared with controls. Single unit recording was performed in the VTA slice. (A) There is no significant difference in spike numbers in the base between controls and EGF-treated rats. (U = 62.0, P = 0.16 by Mann-Whitney’s U-test) (B) For the stimulation, Krebs solution containing NMDA (20 μM) was perfused. The response to NMDA was smaller in EGF-treated rats than in controls (N = 14 cells for controls; N = 13 cells for EGF-treated rats; Animal: N = 5–7).

It is believed that dopamine neural activity in the resting-state results in sustained dopamine release (tonic release) [21]. In agreement, I have previously shown that alterations of the mean firing rate of VTA dopamine neurons (under anesthetized conditions) by chemogenetics are consistent with changes in dopamine levels in the prefrontal cortexes of awake EGF-treated rats and normal rats in the resting-state [13,27]. On the other hand, I reported previously that there was no significant difference in the spike within the burst (SWB), which is a burst firing index, of VTA dopamine neurons between controls and EGF=treated rats under anesthetized conditions [27]. Therefore, the enhancement of tonic release in the prefrontal cortex of EGF-treated rats is assumed to be mainly derived from the increase in the mean firing rate of VTA dopamine neurons in the resting-state, rather than their firing pattern. Then, what mechanisms underlie the elevation of the mean firing rate of VTA dopaminergic neurons in EGF rats? I previously reported that chronic administration of the antipsychotic risperidone ameliorated both the abnormal mean firing rate of VTA dopaminergic neurons and the increased tonic release in the prefrontal cortex [27]. As risperidone primarily exerts antagonistic effects on 5HT2 receptors and dopamine D2 receptors, the abnormal firing of VTA dopamine neurons in EGF-treated rats may be related to these two receptors. To test this assumption, we examined the effects of the chronic coadministration of 5HT2 receptor and D2 receptor antagonists or the single administration of each antagonist on VTA dopaminergic firing (Figure 4A). In agreement with my previous results, I found that the mean firing rate of VTA dopaminergic neurons under anesthesia was enhanced in EGF-treated rats [Kruskal–Wallis test: H = 27.2, P < 0.0001; Scheffe’s comparison: P = 0.0043] (Figure 4B). The chronic coadministration of ritanserin, a 5HT2 receptor antagonist, and haloperidol, a D2 receptor antagonist, ameliorated abnormal mean firing rate of VTA dopaminergic in EGF-treated rats [P = 0.022 by Scheffe’s comparison] as risperidone did (Figure 4B). Moreover, the administration of ritanserin alone demonstrated similar results [P = 0.033 by Scheffe’s comparison]. In contrast, the administration of haloperidol alone exerted no significant effect [P > 0.999 by Scheffe’s comparison] (Figure 4B). In controls, the any drug treatments did not produce significant effects on mean firing rates of VTA dopamine neurons [Kruskal-Wallis test: H = 5.84, P = 0.12] (Figure. 4C). This result indicates that the effect of ritanserin on the activity of VTA dopamine neurons is specific in EGF-treated rats. Our previous study showed that serotoninergic firing in the dorsal raphe nucleus (DRN) was not altered in EGF-treated rats [27]. These results suggest that the abnormal activity of VTA dopaminergic neurons in EGF-treated rats is related to postsynaptic 5HT2 receptor function rather than alteration in serotonin input. To explore the involvement of 5HT2 receptors, we examined how the local administration of a 5HT2 receptor antagonist to the recording site influences dopamine neural activity during recording (Figure 5A). We found that the local administration of ritanserin by iontophoresis increased the dopamine activity in control rats to a greater extent than that in EGF-treated rats [repeated ANOVA: F(1,66) = 8.60, P = 0.0046] (Figure 5B). Next, we applied a selective 5HT2A receptor antagonist, MDL100907, to investigate receptor selectivity and observed a similar effect as that exerted by ritanserin administration [repeated ANOVA: F(1,23) = 4.72, P = 0.041] (Figure 5C). These data suggest that the inhibitory regulation of VTA dopaminergic neurons by endogenous serotonin via 5HT2 receptors, especially 5HT2A receptors, is impaired in EGF-treated rats.

**Figure 4.**
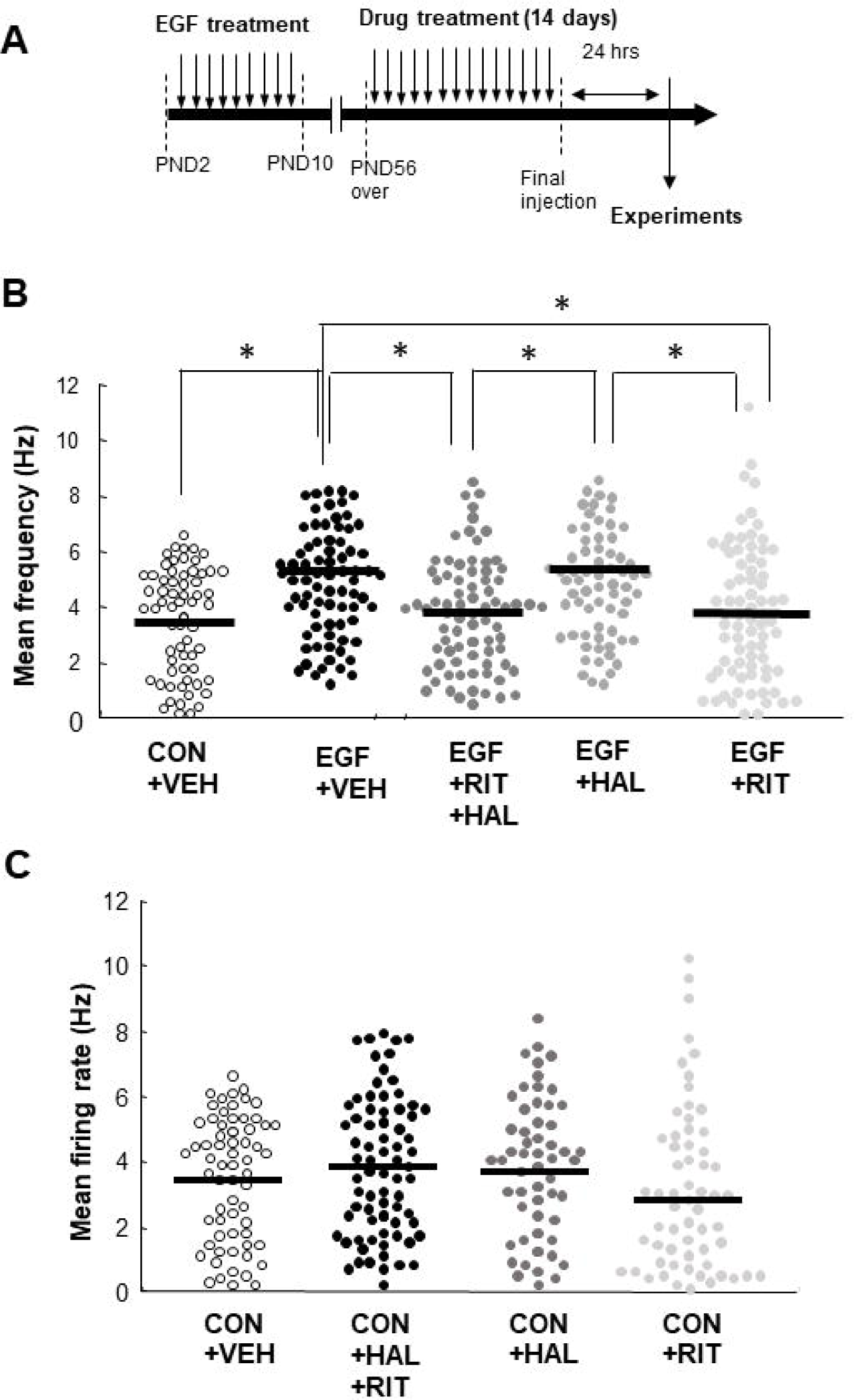
Effect of antagonists against 5HT2 receptors or dopamine D2 receptors on the activity of VTA dopaminergic neurons under anesthetized condition. (A) Experimental procedure. Vehicle (VEH), Ritanserin (RIT), haloperidol (HAL), or combination of ritanserin and haloperidol was administered daily to adult control (CON) and EGF-treated rats (EGF) [over postnatal day (PND) 56] for 2 weeks. In vivo single unit recording was performed 24 h after the final injection of each drug. (B) Ritanserin and coadministration of ritanserin and haloperidol, but not haloperidol, ameliorated the abnormal activity of VTA dopamine neurons in EGF-treated rats (N = 65 cells for CON+VEH; N = 79 cells for EGF+VEH; N = 82 cells for EGF+RIT+HAL; N = 71 cells for EGF+HAL; N = 82 cells for EGF+RIT; Animal: N = 5–7). * *P* < 0.05. (C) All treatments did not affect the activity of VTA dopamine neurons in controls (N = 65 cells for CON+VEH; N = 77 cells for CON+RIT+HAL; N = 61 cells for CON+HAL; N = 64 cells for CON+RIT; Animal: N = 5–7).

**Figure 5.**
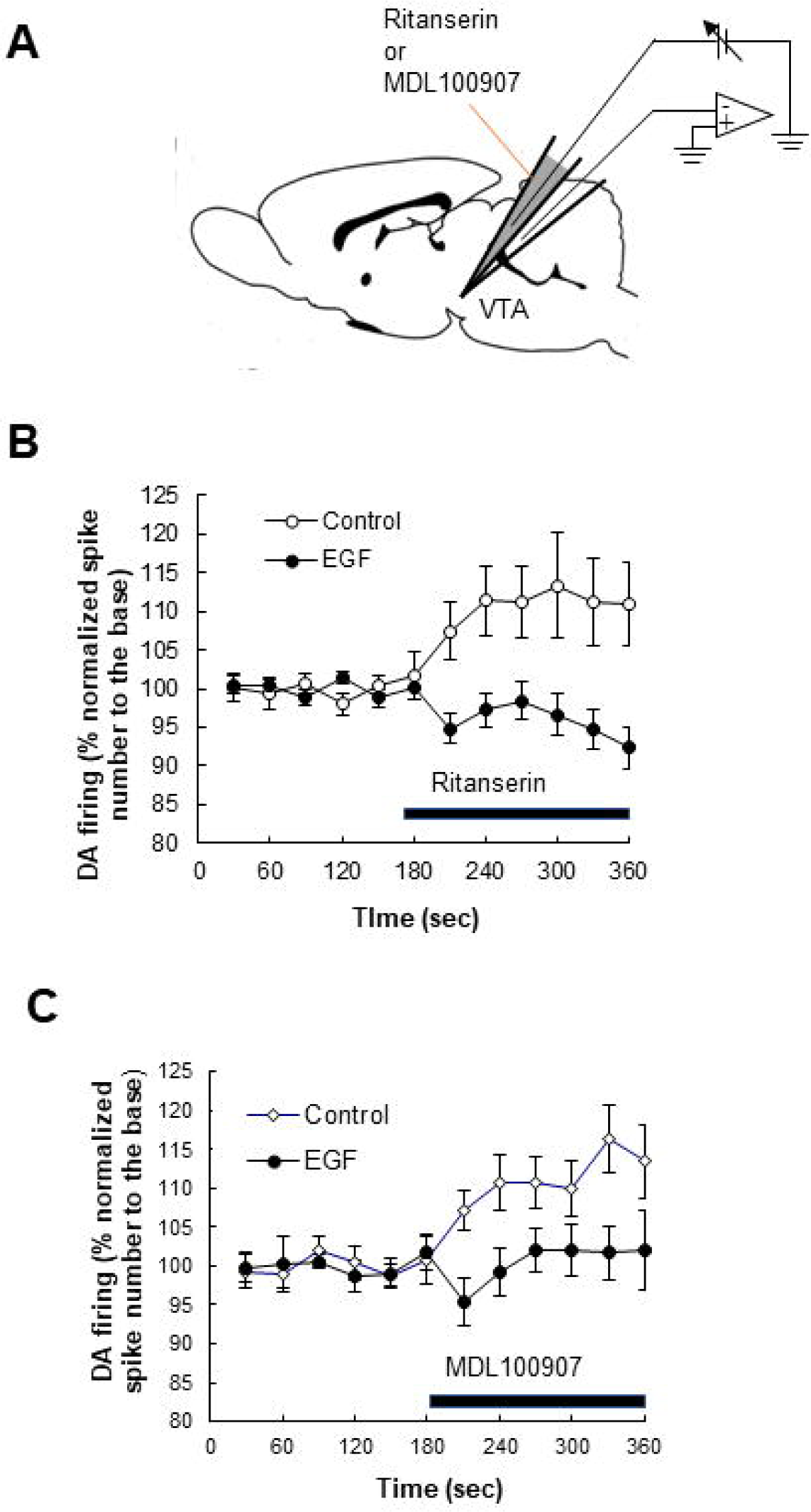
Effects of local administration of 5HT2 receptor antagonists on the activity of VTA dopaminergic neurons in vivo. (A) Single unit recording was performed in the VTA. (B) 5HT2 receptor antagonist ritanserin was introduced into the recording site (N = 34 cells for controls; N = 32 cells for EGF-treated rats; Animal: N = 8–9). (C) The selective 5HT2A4 receptor antagonist MDL100907 was introduced into the recording site (N = 13 cells for controls; N = 10 cells for EGF-treated rats; Animal: N = 3–4).

The 5HT2A receptor is coupled to the G-protein Gq [36,37]. Activation of this receptor enhances neuronal excitability and activity [38–41]. Therefore, the inhibitory effect of 5HT2A receptors observed in this study raises the possibility that serotonin acts indirectly on VTA dopamine neurons via GABA, rather than acting directly on VTA dopaminergic neurons. To confirm this possibility, we examined the effect of serotonin on VTA dopaminergic activity in the presence and absence of local GABA inputs using VTA slices. In the recordings in normal artificial cerebrospinal fluid (ACSF), I found response of VTA dopamine neurons to serotonin contains more repressive component in controls compared with EGF-treated rats [repeated ANOVA: F(2,87) = 7.12, P = 0.0014; Fisher’s LSD comparison: P = 0.0006 (Control vs EGF)] (Figure.6B). This repressive effect in controls shifted toward less repressive in the presence of ritanserin [repeated ANOVA: F(1,12) = 6.37, P = 0.027] (Figure. 6A), suggesting that the repressive component in the response of VTA dopamine neurons to serotonin is mediated by 5HT2 receptors. In the presence of the GABA-A receptor antagonist bicuculline, the repressive component in the serotoninergic response of VTA dopamine neurons in controls is reduced, resulting in abolishment of a significant difference in the serotoninergic responses between controls and EGF-treated rats [repeated ANOVA: F(2,50) = 2.38, P = 0.103; Fisher’s LSD comparison: P = 0.70 (Control vs EGF)]. This result indicates that serotonin normally increases GABAergic inputs to VTA dopamine neurons via 5HT2 receptors and indirectly inhibits dopaminergic neurons and that this serotonergic regulation is disrupted in EGF-treated rats. These data suggest that the enhanced tonic release in the prefrontal cortex of EGF-treated rats is originated from this disinhibition of VTA dopaminergic neurons. Interestingly, the chronic administration of risperidone normalized the serotonergic responsiveness of VTA dopaminergic neurons in EGF-treated rats [Fisher’s LSD comparison: P = 0.025 (EGF vs. EGF+RIS)], but not in the presence of bicuculline [Fisher’s LSD comparison: P = 0.102 (EGF vs. EGF+RIS)] (Figures 6C and D). Hence, the mechanism underlying the action of chronic risperidone administration involves the restoration of the indirect serotonergic inhibition of VTA dopaminergic neurons via GABA.

**Figure 6.**
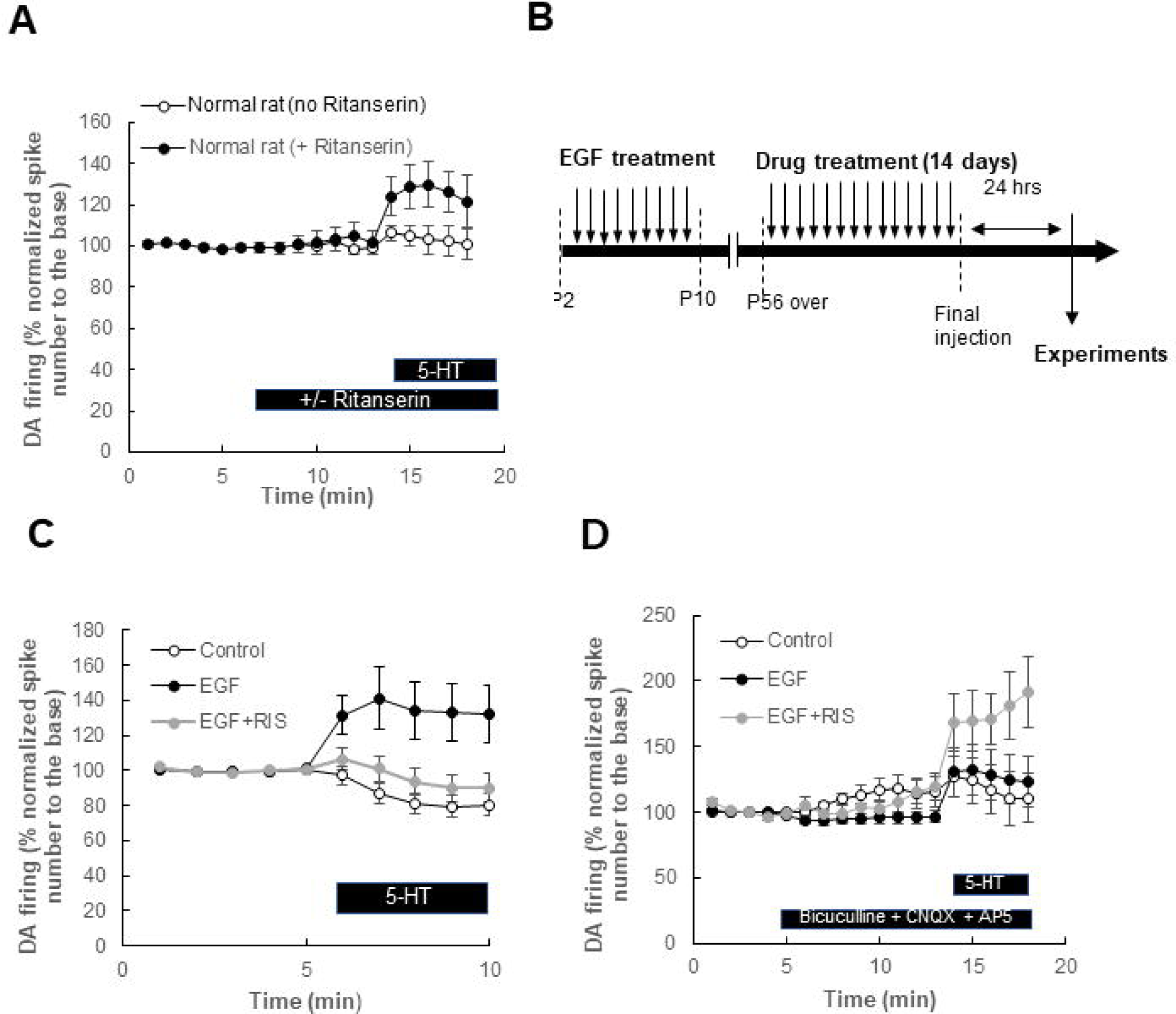
Influence of serotonin on the activity of VTA dopaminergic neurons is affected by local GABA neurons. (A) Effect of 5HT2 receptor blockade on the response of VTA dopaminergic neurons to serotonin. Single unit recording of a dopaminergic neuron was performed using the VTA slices of control rats in the presence of ritanserin (10 µM). (N = 7 cells for each) (B) Experimental procedure. (C) The alterations of the normalized spike number of VTA dopaminergic neurons to the base after serotonin application was investigated in the absence of GABAergic and glutamatergic inhibitors. (Control: N = 29 cells; EGF: N = 33 cells; EGF + Risperidone (RIS): N = 28 cells; Animal: N = 7–12) (D) investigated in the presence of GABAergic and glutamatergic inhibitors (Control: N = 22 cells; EGF: N = 17 cells; EGF + Risperidone (RIS): N = 14 cells; Animal: N = 4–5).

To elucidate serotonergic regulation of VTA dopamine neurons and its role in the regulation of social behavior, the effects of chronic activation of serotonergic neurons in the DRN by chemogenetics on VTA dopamine neuron activity and social behavior were examined in normal rats (Figure. 7A). I found that serotoninergic neurons in the DRN positively regulates social behavior and also interacts with VTA dopaminergic neurons. Chronic activation of serotonergic neurons in the DRN by chemogenetics (Mann-Whitney’s U-test: P = 0.004) reduces VTA dopaminergic activity in the resting state (Mann-Whitney’s U-test: P = 0.035) and increases social interaction (Figures. 7B, C and D). Because the attenuation of VTA dopamine neuronal activity in the resting state causally increases social interaction in normal rats as well [13], these results indicate that serotonin regulates social behavior through VTA dopamine neurons. In this experiment, a significant correlation was observed between the averages of the firing activity of VTA dopamine neurons in the individual animals and social interaction (Pearson’s correlation: R = -0.622, P = 0.027) (Figure. 7F). In contrast, the averages of serotonergic neuronal activities in the DRN of individual animals exhibited no significant correlation with social interaction, although activation of serotonergic neurons has causality with changes in social interaction (Pearson’s correlation: R = 0.38, P = 0.29) (Figure 7E). This result suggests that only a subset of DRN serotonergic neurons is involved in social behavior regulation.

**Figure 7.**
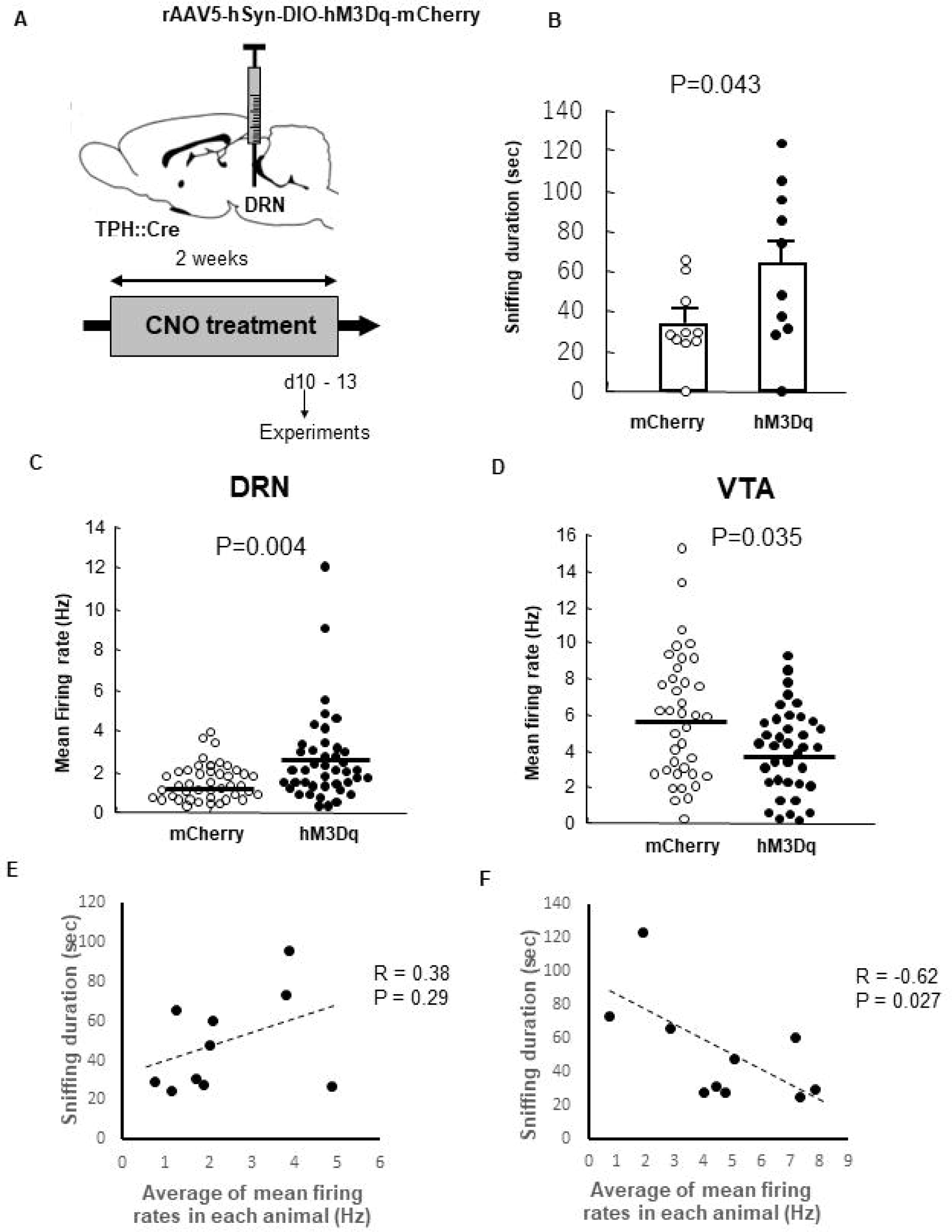
Chronic chemogenetic activation of serotonergic neurons in the dorsal raphe nucleus promotes social behavior via the repression of sustained activity of dopaminergic neurons in the ventral tegmental area. (A) rAAV5-hsyn-DIO-hM3Dq-mCherry or rAAV5-hsyn-DIO-mCherry was infused into the dorsal raphe of male knockin rats expressing Cre recombinase by a tryptophan hydroxylase 2 gene promoter driving (TPH::Cre). Clozapine-N-oxide (CNO) was administered continuously from a mini-osmotic pump implanted under the skin for 2 weeks. (B) Social interaction (N = 10 for each). (C) Mean firing rates of serotonergic neurons in the dorsal raphe nucleus (DRN) (mCherry; N = 44 cells; hM3Dq: N = 43 cells; N = 5 animals for each). Mean firing rates of dopaminergic neurons in the ventral tegmental area (VTA) (mCherry; N = 38 cells; hM3Dq: N = 43 cells; N = 5 animals for each). (E) Correlation between social interaction and average of mean firing rates of DRN serotonergic neurons in each animal. (N = 5 animals for each) (F) Correlation between social interaction and average of mean firing rates of VTA dopaminergic neurons in each animal. (N = 5 animals for each). The results in this figure were obtained by reanalysis of our previously acquired data [50].

## Discussion

We investigated the neural mechanisms underlying the abnormal tonic and phasic dopamine releases in an animal model of schizophrenia. The primary findings were that 1) the activation of dopaminergic neurons by NMDA receptors is blunt in EGF-treated rats and 2) the impairment of the indirect inhibition of VTA dopaminergic neurons via GABA by 5HT2A receptors enhances activity of VTA dopamine neurons in the resting state in the model. It has been reported that pharmacological inhibition of NMDA receptors in the VTA and ablation of NMDA receptors in dopaminergic neurons impair phasic dopamine release [42]. These data support the finding of our study that the diminished stimulus-responsiveness of VTA dopaminergic neurons contributes to the attenuation of the phasic dopamine release in the prefrontal cortex of EGF-treated rats. Moreover, another study reported that systemic ablation of the NMDA receptor subunit NR1 exhibits increased tonic activity in addition to reduced phasic firing of dopaminergic neurons [43]. Therefore, it remains to be controversial whether the dysfunction of NMDA receptors in neurons other than dopaminergic neurons may also be involved in the enhancement of tonic dopamine release in EGF-treated rats. Nevertheless, the lack of alteration in the expression of NR1 in VTA tissues of EGF-treated rats (data not shown) implies that the hypo-NMDA receptor function in these rats is fundamentally different from that in the NR1 knock-down mice.

The increased GABAergic input from the lateral globus pallidus (GP) to the substantia nigra pars reticulata (SNr) is observed in EGF-treated rats [33]. We have thought that this enhanced inhibitory input might reduce the activity of SNr GABA neurons that suppress mid-brain dopamine neurons, resulting in disinhibition of dopamine neurons. However, as GP hyperactivity is originally caused by excessive innervation from dopamine neurons in the substantia nigra pars compacta (SNc) [29], which of the dopaminergic neurons or GP GABA neurons causes an abnormality in the basal ganglia circuit remains unclear. The finding that serotonergic regulation of inhibitory inputs to VTA dopamine neurons is impaired suggests that dopamine may be responsible for the basal ganglia circuit abnormalities in EGF-treated rats. 5HT2 receptor-induced depolarization of SNr GABA neurons is restricted by activating the GABA-B receptor [58,59]. Therefore, enhanced GABAergic inputs from the GP to the SNr are expectedly involved in the attenuation of 5-HT2 receptor-mediated signaling in the SNr neurons as well. However, given that the altered serotonergic regulation of VTA dopamine neurons remained even in slices with cutoff of inhibitory inputs from the GP, the insight that 5-HT2 receptor-induced regulation itself may be impaired by some factors is more rational. However, it is of note that the inhibitory inputs themselves to dopamine neurons may be also reduced in EGF-treated rats since there is a trend that bicuculine itself slightly increases activities of VTA dopamine neurons in slices of controls compared with EGF-treated rats (Figure. 6D) (P = 0.059 by Fisher’s LSD). On the other hand, this trend is not observed between EGF-treated rats and EGF-treated rats with the risperidone treatment (Figure. 6D), suggesting the consequence of this antipsychotic treatment may not be substantially equivalent to restoration of normal neural circuits.

Both the excitatory and inhibitory regulations of VTA dopamine neurons are impaired in EGF-treated rats. However, the result of the basal firing of VTA dopamine neurons is higher in EGF-treated rats than in controls, suggesting the effects of disinhibition exceed those of hypoexcitation. Why, then, did this disinhibition effect not favor phasic release? This issue may be related to the differential effects of tonic GABAergic inputs on the firing dopamine neurons depending on their firing patterns. The single spike (regular firing) of dopamine neurons has been reported to be sensitive to tonic GABAergic inputs, whereas the NMDA-induced burst firing is less affected [60]. Given that no significant difference was found in the burst firing of VTA dopamine neurons between EGF-treated rats and controls under anesthesia [27], this difference in inhibitory effect on dopamine neurons between the controls and EGF-treated rats may not be significantly sufficient to produce the superiority of EGF-treated rats in burst firing rates compared to controls. Indeed, in vivo recordings of VTA dopamine neurons under awake conditions revealed that burst firing in controls increased to the same level as in EGF-treated rats during social stimulation [27]. Therefore, the disinhibition of dopaminergic neurons in EGF rats may not contribute to the induction of a larger phasic release than controls.

In the present study, we demonstrated that the effect of an atypical antipsychotic drug is attributed to restoring the serotonergic indirect inhibition of VTA dopaminergic neurons via GABA. We have previously reported that this abnormal VTA dopaminergic activity has causality to attenuate social interaction in EGF-treated rats [27]. Therefore, it is reasonable to consider that this restoration of the serotonergic inhibition is related to the mechanism underlying the improvement of social deficit in EGF-treated rats by atypical antipsychotics. Interestingly, a typical antipsychotic drug, haloperidol, with low affinity against 5HT2 receptors did not exhibit this efficacy. The dose of haloperidol used in the present study was twice the dose that improves the hyperactivity of EGF-treated rats [23]. Considering this viewpoint, it is believed that the antagonism of D2 receptors alone, rather than simply a matter of dosage, cannot ameliorate this abnormal firing of dopaminergic neurons. This difference in the effects of these drugs on VTA dopaminergic neurons may be useful to explain the statistically significant efficacy of atypical antipsychotics on the negative symptoms of schizophrenia compared with typical antipsychotics [44,45].

Previous studies have reported that 5HT2A receptors exhibit high expression in the cerebral cortex [46–48]. Based on this localization of 5HT2A receptors, it has been thought that atypical antipsychotics may exert therapeutic effects by significantly affecting the cerebral cortex [49]. The effect on VTA dopamine neurons observed in the present study may be out of the general interpretation for mechanisms of drug efficacy against social deficits. However, the contribution of cortical serotonin to regulation of social behavior remains to be elucidated. The present study shows that a subpopulation of DRN serotonin neurons involved in the regulation of VTA dopamine neurons contribute to control of social behavior. In other words, the serotonergic regulation of the ventral tegmental area may be far more important for social behavior control than serotonergic control over other brain regions. Considering these results, the improvement of impaired serotonergic regulation in the VTA is important for efficacy of atypical antipsychotics against social deficits.

In conclusion, the present study suggests that diminished serotoninergic regulation of VTA dopamine neurons via GABA neurons and hypofunction of NMDA receptors in VTA dopamine neuron are independently associated with abnormalities in tonic and phasic dopamine releases in the prefrontal cortex of a schizophrenia-like model. In particular, this mechanism underlying the abnormal tonic dopamine release may be strongly involved in the mechanism underlying the action of atypical antipsychotic drugs, which has not yet been understood in detail. To the best of our knowledge, this study is the first report to provide a clear insight into the roles of the three neurotransmitters in the mechanism underlying the significant efficacy of atypical antipsychotic drugs compared with typical antipsychotic drugs in chronic drug treatments. However, it should be of note that it may be an oversimplification to explain abnormalities in dopamine releases under awake conditions by only abnormal firing activity of dopamine neurons. Firing activity of VTA dopamine neurons under awake condition in my previous study is not completely consistent with the dynamics of the dopamine release in the prefrontal cortex in this study [27]. Given the present results support the alterations of activity of VTA dopamine neurons in awake EGF-treated rats, these are considered components involved in the neural mechanisms underlying the abnormal dopamine releases; however, the existence of other components cannot be ruled out.

## Acknowledgement

I thank Prof. Hiroyuki Nawa, Wakayama Medical University, for his valuable comments for this study. I thank Dr. Hisaaki Namba, Wakayama Medical University, for his informative advices for electrophysiology and Ms. Eiko Kitayama, Niigata University, for her careful animal care. I also thank Dr. Nobuyuki Takei, Niigata University, and Prof. Tomoo Homma, Maebashi Institute of Technology, for their valuable comments for this manuscript.

## Data and materials availability

All the data required to support the conclusion of this paper are presented within the paper and its supplemental materials. The data that support the findings of this study are available from the corresponding author upon reasonable request.

## References

1. Chaudhury D, Walsh JJ, Friedman AK, Juarez B, Ku SM, Koo JW, Ferguson D, Tsai HC, Pomeranz L, Christoffel DJ, Nectow AR, Ekstrand M, Domingos A, Mazei-Robison MS, Mouzon E, Lobo MK, Neve RL, Friedman JM, Russo SJ, Deisseroth K, Nestler EJ & Han MH (2013) Rapid regulation of depression-related behaviours by control of midbrain dopamine neurons. Nature 493, 532–536.

2. Gunaydin LA, Grosenick L, Finkelstein JC, Kauvar IV, Fenno LE, Adhikari A, Lammel S, Mirzabekov JJ, Airan RD, Zalocusky KA, Tye KM, Anikeeva P, Malenka RC & Deisseroth K (2014) Natural neural projection dynamics underlying social behavior. Cell 157(7), 1535–1551.

3. Sawaguchi T & Goldman-Rakic PS (1991) D1 dopamine receptors in prefrontal cortex: involvement in working memory. Science 251(4996), 947–950.

4. Simon H, Scatton B & Moal ML (1980) Dopaminergic A10 neurones are involved in cognitive functions. Nature 286(5769), 150–151.

5. Mohebi A, Pettibone JR, Hamid AA, Wong JT, Vinson LT, Patriarchi T, Tian L, Kennedy RT & Berke JD (2019) Dissociable dopamine dynamics for learning and motivation. Nature 570(7759), 65–70.

6. Davis KL, Kahn RS, Ko G & Davidson M (1991) Dopamine in schizophrenia: a review and reconceptualization. Am J Psychiatry 148, 1474–1486.

7. Arnsten AF, Girgis RR, Gray DL & Mailman RB (2017) Novel Dopamine Therapeutics for Cognitive Deficits in Schizophrenia. Biol Psychiatry 81, 67–77.

8. Dolan RJ, Fletcher P, Frith CD, Friston KJ, Frackowiak RS & Grasby PM (1995) Dopaminergic modulation of impaired cognitive activation in the anterior cingulate cortex in schizophrenia. Nature 378(6553), 180–182.

9. Jentsch JD, Redmond DE Jr, Elsworth JD, Taylor JR, Youngren KD & Roth RH (1997) Enduring cognitive deficits and cortical dopamine dysfunction in monkeys after long-term administration of phencyclidine. Science 277(5328), 953–955.

10. Takahashi K, Nakagawasai O, Sakuma W, Nemoto W, Odaira T, Lin JR, Onogi H, Srivastava LK & Tan-No K (2019) Prenatal treatment with methylazoxymethanol acetate as a neurodevelopmental disruption model of schizophrenia in mice. Neuropharmacology 150, 1–14.

11. Didriksen M, Fejgin K, Nilsson SRO, Birknow MR, Grayton HM, P. Larsen PH, Lauridsen JB, Nielsen V, Celada P, Santana N, Kallunki P, Christensen KV, Werge TM, Stensbøl TB, Egebjerg J, Gastambide F, Artigas F, Bastlund JF & Nielsen J (2017) Persistent gating deficit and increased sensitivity to NMDA receptor antagonism after puberty in a new mouse model of the human 22q11.2 microdeletion syndrome: a study in male mice. J Psychiatry Neurosci 42(1), 48–58.

12. Winter C, Djoda-Irani A, Sohr R, Morgenstern R, Feldon J, Juckel G & Meyer U (2009) Prenatal immune activation leads to multiple changes in basal neurotransmitter levels in the adult brain: implications for brain disorders of neurodevelopmental origin such as schizophrenia. Int J Neuropsychopharmacol 12(4), 513–524.

13. Sotoyama H, Inaba H, Iwakura Y, Namba H, Takei N, Sasaoka T & Nawa H (2022) The dual role of dopamine in the modulation of information processing in the prefrontal cortex underlying social behavior. FASEB J 36(2), e22160.

14. Sulzer D, Cragg SJ & Rice ME (2016) Striatal dopamine neurotransmission: regulation of release and uptake. Basal Ganglia 6(3), 123–148.

15. Ojeil N, Moore H, D’Souza D, Malison RT, Huang Y, Lim K, Nabulsi N, Carson RE, Lieberman JA & Abi-Dargham A (2015) Deficits in prefrontal cortical and extrastriatal dopamine release in schizophrenia: a positron emission tomographic functional magnetic resonance imaging study. JAMA Psychiatry 72(4), 316–324.

16. Frankle WG, Himes M, Mason NS, Mathis CA & Narendran R (2022) Prefrontal and Striatal Dopamine Release Are Inversely Correlated in Schizophrenia. Biol Psychiatry 92(10), 791–799.

17. Olsson SK, Andersson AS, Linderholm KR, Holtze M, Nilsson-Todd LK, Schwieler L, Olsson E, Larsson K, Engberg G & Erhardt S (2009) Elevated levels of kynurenic acid change the dopaminergic response to amphetamine: implications for schizophrenia. Int J Neuropsychopharmacol 12(4), 501–512.

18. Flagstad P, Mørk A, Glenthøj BY, van Beek J, Michael-Titus AT & Didriksen M (2004) Disruption of neurogenesis on gestational day 17 in the rat causes behavioral changes relevant to positive and negative schizophrenia symptoms and alters amphetamine-induced dopamine release in nucleus accumbens. Neuropsychopharmacology 29(11), 2052–2064.

19. Jones SR, Gainetdinov RR, Wightman RM & Caron MG (1998) Mechanisms of amphetamine action revealed in mice lacking the dopamine transporter. J Neurosci 18(6), 1979–1986.

20. Floresco SB, West AR, Ash B, Moore H & Grace AA (2003) Afferent modulation of dopamine neuron firing differentially regulates tonic and phasic dopamine transmission. Nat Neurosci 6(9), 968–973.

21. Grace AA (1991) Phasic versus tonic dopamine release and the modulation of dopamine system responsivity: a hypothesis for the etiology of schizophrenia. Neuroscience 41(1), 1–24.

22. Rubio R, Aguilar R, Bustamante M, Muñoz E, Vázquez-Santiago M, Santano R, Vidal M, Melero NR, Parras D, Serra P, Santamaria P, Carolis C, Izquierdo L, Gómez-Roig MD, Dobaño C, Moncunill G & Mazarico E (2022) Maternal and neonatal immune response to SARS-CoV-2, IgG transplacental transfe r and cytokine profile. Front Immunol 13, 999136.

23. Futamura T, Kakita A, Tohmi M, Sotoyama H, Takahashi H & Nawa H (2003) Neonatal perturbation of neurotrophic signaling results in abnormal sensorimotor gating and social interaction in adults: implication for epidermal growth factor in cognitive development. Mol Psychiatry 8(1), 19–29.

24. Futamura T, Toyooka K, Iritani S, Niizato K, Nakamura R, Tsuchiya K, Someya T, Kakita A, Takahashi H & Nawa H (2002) Abnormal expression of epidermal growth factor and its receptor in the forebrain and serum of schizophrenic patients. Mol Psychiatry 7(7), 673–682.

25. Mizuno M, Sotoyama H, Namba H, Shibuya M, Eda T, Wang R, Okubo T, Nagata K, Iwakura Y & Nawa H (2013) ErbB inhibitors ameliorate behavioral impairments of an animal model for schizophrenia: implication of their dopamine-modulatory actions. Transl Psychiatry 3(4), e252.

26. Jodo E, Inaba H, Narihara I, Sotoyama H, Kitayama E, Yabe H, Namba H, Eifuku S & Nawa H (2019) Neonatal exposure to an inflammatory cytokine, epidermal growth factor, results in the deficits of mismatch negativity in rats. Sci Rep 9(1), 7503.

27. Sotoyama H, Namba H, Kobayashi Y, Hasegawa T, Watanabe D, Nakatsukasa E, Sakimura K, Furuyashiki T & Nawa H (2021) Resting-state dopaminergic cell firing in the ventral tegmental area negatively regulates affiliative social interactions in a developmental animal model of schizophrenia. Transl Psychiatry 11(1), 236.

28. Paxinos P & Watson C. (1998) The Rat Brain in Stereotaxic Coordinates (4^th^ edition). Academic Press, San Diego, CA, USA.

29. Sotoyama H, Zheng Y, Iwakura Y, Mizuno M, Aizawa M, Shcherbakova K, Wang R, Namba H & Nawa H (2022) Pallidal hyperdopaminergic innervation underlying D2 receptor-dependent behavioral deficits in the schizophrenia animal model established by EGF. PLoS One 6(10), e25831.

30. Mameli-Engvall M, Evrard A, Pons S, Maskos U, Svensson TH, Changeux JP & Faure P (2006) Hierarchical control of dopamine neuron-firing patterns by nicotinic receptors. Neuron 50(6), 911–921.

31. Lammel S, Hetzel A, Häckel O, Jones I, Liss B, Roeper J. (2008) Unique properties of mesoprefrontal neurons within a dual mesocorticolimbic dopamine system. Neuron. 57(5):760–773.

32. Rainer Q, Speziali S, Rubino T, Dominguez-Lopez S, Bambico FR, Gobbi G & Parolaro D (2014) Chronic nandrolone decanoate exposure during adolescence affects emotional behavior and monoaminergic neurotransmission in adulthood. Neuropharmacology 83, 79–88.

33. Sotoyama H, Namba H, Chiken S, Nambu A & Nawa H (2013) Exposure to the cytokine EGF leads to abnormal hyperactivity of pallidal GABA neurons: implications for schizophrenia and its modeling. J Neurochem 126(4), 518–528.

34. Takahata R & Moghaddam B (1998) Glutamatergic regulation of basal and stimulus-activated dopamine release in the prefrontal cortex. J Neurochem 71(4), 1443–1449.

35. Wang B, You ZB & Wise RA (2012) Heroin self-administration experience establishes control of ventral tegmental glutamate release by stress and environmental stimuli. Neuropsychopharmacology 37(13), 2863–2869.

36. Maroteaux L, Ayme-Dietrich E, Aubertin-Kirch G, Banas S, Quentin E, Lawson R & Monassier L (2017) New therapeutic opportunities for 5-HT_2_ receptor ligands. Pharmacol Ther 170, 14–36.

37. Odagaki Y & Toyoshima R (2013) Activation of Gq proteins coupled with 5-HT2 receptors in rat cerebral cortical membranes assessed by antibody-capture scintillation proximity assay/[S]GTPγS binding. Pharmacology 92(1-2), 2–10.

38. Larson EA, Metzen MG & Chacron MJ (2014) Serotonin modulates electrosensory processing and behavior via 5-HT2-like receptors. Neuroscience 271, 108–118.

39. Zhang ZW (2003) Serotonin induces tonic firing in layer V pyramidal neurons of rat prefrontal cortex during postnatal development. J Neurosci 23(8), 3373–3384.

40. Araneda R & Andrade R (1991) 5-Hydroxytryptamine-2 and 5-hydroxytryptamine-1A receptors mediate opposing responses on membrane excitability in rat association cortex. Neuroscience 40, 399–412.

41. Amargós-Bosch M, Bortolozzi A, Puig MV, Serrats J, Adell A, Celada P, Toth M, Mengod G & Artigas F (2004) Co-expression and in vivo interaction of serotonin1A and serotonin2A receptors in pyramidal neurons of prefrontal cortex. Cereb Cortex 14, 281–299.

42. Parker JG, Zweifel LS, Clark JJ, Evans SB, Phillips PE & Palmiter RD (2010) Absence of NMDA receptors in dopamine neurons attenuates dopamine release but not conditioned approach during Pavlovian conditioning. Proc Natl Acad Sci U S A 107(30), 13491–13496.

43. Ferris MJ, Milenkovic M, Liu S, Mielnik CA, Beerepoot P, John CE, España RA, Sotnikova TD, Gainetdinov RR, Borgland SL, Jones SR & Ramsey AJ (2014) Sustained N-methyl-d-aspartate receptor hypofunction remodels the dopamine system and impairs phasic signaling. Eur J Neurosci 40(1), 2255–2263.

44. Leucht S, Corves C, Arbter D, Engel RR, Li C & Davis JM (2009) Second-generation versus first-generation antipsychotic drugs for schizophrenia: a meta-analysis. Lancet 373(9657), 31–41.

45. Zhang JP, Gallego JA, Robinson DG, Malhotra AK, Kane JM & Correll CU (2013) Efficacy and safety of individual second-generation vs. first-generation antipsychotics in first-episode psychosis: a systematic review and meta-analysis. Int J Neuropsychopharmacol 16(6), 1205–1218.

46. López-Giménez JF, Vilaró MT, Palacios JM & Mengod G (2001) Mapping of 5-HT2A receptors and their mRNA in monkey brain: [3H]MDL100,907 autoradiography and in situ hybridization studies. J Comp Neurol 429(4), 571–589.

47. Cornea-Hébert V, Riad M, Wu C, Singh SK & Descarries L (1999) Cellular and subcellular distribution of the serotonin 5-HT2A receptor in the central nervous system of adult rat. J Comp Neurol 409(2), 187–209.

48. Pompeiano M, Palacios JM & Mengod G (1994) Distribution of the serotonin 5-HT2 receptor family mRNAs: comparison between 5-HT2A and 5-HT2C receptors. Brain Res Mol Brain Res 23(1-2), 163–178.

49. Bortolozzi A, Díaz-Mataix L, Scorza MC, Celada P & Artigas F (2005) The activation of 5-HT receptors in prefrontal cortex enhances dopaminergic activity. J Neurochem 95(6), 1597–607.

50. Nawa H, Kikawa M, Namba H & Sotoyama H (2021) Mannsei Sutoresu Fuka to Kouutsuzai ni yoru Nounai Monoaminn Shinnkei Katsudou no Barannsu Seigyo (Regulation of the balance among monoamine neural activities in the brain by chronic stress application and antidepressant). SEITAI NO KAGAKU 72(5), 497-500.

51. Yamashita M, Chattopadhyay S, Fensterl V, Saikia P, Wetzel J,; Sen GC (2012) Epidermal growth factor receptor is essential for Toll-like receptor 3 signaling. Sci. Signal. 5, ra50.

52. Singh AB, Harris RC (2005) Autocrine, paracrine and juxtacrine signaling by EGFR ligands. Cell. Signal. 17, 1183–1193

53. Stenkrona P, Matheson GJ, Halldin C, Cervenka S & Farde L (2019) D1-Dopamine Receptor Availability in First-Episode Neuroleptic Naive Psychosis Patients. Int J Neuropsychopharmacol 22(7), 415–425.

54. Okubo Y, Suhara T, Suzuki K, Kobayashi K, Inoue O, Terasaki O, Someya Y, Sassa T, Sudo Y, Matsushima E, Iyo M, Tateno Y & Toru M (1997) Decreased prefrontal dopamine D1 receptors in schizophrenia revealed by PET. Nature 385(6617), 634–636.

55. Lehrer DS, Christian BT, Kirbas C, Chiang M, Sidhu S, Short H, Wang B, Shi B, Chu KW, Merrill B & Buchsbaum MS (2010) 18F-fallypride binding potential in patients with schizophrenia compared to healthy controls. Schizophr Res 122(1-3), 43–52.

56. Lindström LH, Gefvert O, Hagberg G, Lundberg T, Bergström M, Hartvig P & Långström B (1999) Increased dopamine synthesis rate in medial prefrontal cortex and striatum in schizophrenia indicated by L-(beta-11C) DOPA and PET. Biol Psychiatry 46(5), 681–688.

57. Rao N, Northoff G, Tagore A, Rusjan P, Kenk M, Wilson A, Houle S, Strafella A, Remington G & Mizrahi R (2019) Impaired Prefrontal Cortical Dopamine Release in Schizophrenia During a Cognitive Task: A [11C]FLB 457 Positron Emission Tomography Study. Schizophr Bull 45(3), 670–679.

58. Invernizzi RW, Pierucci M, Calcagno E, Di Giovanni G, Di Matteo V, Benigno A, Esposito E. (2007 Selective activation of 5-HT(2C) receptors stimulates GABA-ergic function in the rat substantia nigra pars reticulata: a combined in vivo electrophysiological and neurochemical study. Neuroscience. 144(4), 1523–1535.

59. Stanford IM, Lacey MG. (1996) Differential actions of serotonin, mediated by 5-HT1B and 5-HT2C receptors, on GABA-mediated synaptic input to rat substantia nigra pars reticulata neurons in vitro. J Neurosci. 16(23), 7566–7573.

60. Lobb CJ, Wilson CJ, Paladini CA. (2010) A dynamic role for GABA receptors on the firing pattern of midbrain dopaminergic neurons. J Neurophysiol. 104(1), 403–413.

61. Sotoyama H, Namba H, Tohmi M, Nawa H. (2023) Schizophrenia Animal Modeling with Epidermal Growth Factor and Its Homologs: Their Connections to the Inflammatory Pathway and the Dopamine System. Biomolecules. 13(2):372.

62. Fukata M, Chen A, Vamadevan AS, Cohen J, Breglio K, Krishnareddy S, Hsu D, Xu R, Harpaz N, Dannenberg AJ. et al. (2007) Toll-like receptor-4 promotes the development of colitis-associated colorectal tumors. Gastroenterology 133, 1869–1881.

63. Damiano V, Caputo R, Bianco R, D’Armiento FP, Leonardi A, De Placido S, Bianco AR, Agrawal S, Ciardiello F, Tortora G (2006) Novel toll-like receptor 9 agonist induces epidermal growth factor receptor (EGFR) inhibition and synergistic antitumor activity with EGFR inhibitors. Clin. Cancer Res. 12, 577–583.

